# Natively Oxidized Amino Acid Residues in the Spinach PS I-LHC I Supercomplex

**DOI:** 10.1101/826362

**Authors:** Ravindra Kale, Larry Sallans, Laurie K. Frankel, Terry M. Bricker

## Abstract

Reactive oxygen species (ROS) production is an unavoidable byproduct of electron transport under aerobic conditions. Photosystem II (PS II), the cytochrome *b*_6_*f* complex and Photosystem I (PS I) are all demonstrated sources of ROS. It has been proposed that PS I produces substantial levels of a variety of ROS including 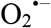 ^1^O_2_, H_2_O_2_ and, possibly, 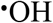, however, the site(s) of ROS production within PS I has been the subject of significant debate. We hypothesize that amino acid residues close to the sites of ROS generation will be more susceptible to oxidative modification than distant residues. In this study, we have identified oxidized amino acid residues in spinach PS I which was isolated from field-grown spinach. The modified residues were identified by high-resolution tandem mass spectrometry. As expected, many of the modified residues lie on the surface of the complex. However, a well-defined group of oxidized residues, both buried and surface-exposed, lead from the chl a’ of P_700_ to the surface of PS I. These residues (PsaB: ^609^F, ^611^E, ^617^M, ^619^W, ^620^L, and PsaF: ^139^L, ^142^A,^143^D) may identify a preferred route for ROS, probably ^1^O_2_, to egress the complex from the vicinity of P_700_. Additionally, two buried residues located in close proximity to A_1B_ (PsaB:^712^H and ^714^S) were modified, which may be consistent with A_1B_ being a source of 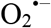. Surprisingly, no oxidatively modified residues were identified in close proximity to the 4Fe-FS clusters F_X_, F_A_ or F_B_. These cofactors had been identified as a principal targets for ROS damage in the photosystem. Finally, a large number of residues located in the hydrophobic cores of Lhca1-Lhca4 are oxidatively modified. These appear to be the result of ^1^O_2_ production by the distal antennae for the photosystem.

## Introduction

In higher plants, the Photosystem I - Light Harvesting Complex I supercomplex (PS ILHC I) acts as a light-energy driven plastocyanin - ferredoxin oxidoreductase. Crystal structures are available for PS I-LHC I from red algae (*Cyanidioschyzon merolae* strain 10D, 4.0 Å (Antoshvili et al. 2019; Pi et al. 2018), green algae (*Chlamydomonas reinhardtii*, 2.9 Å, (Suga et al. 2019)) and angiosperms (*Pisum sativum*, 2.8 Å (Mazor et al. 2017; Qin et al. 2015).

Higher plant PS I-LHC I is a 650 kDa monomeric complex consisting of at least 16 subunits (PS I: 12 core subunits (PsaA-PsaO) and LHC I: 4 subunits (Lhca1-Lhca4)). Numerous prosthetic groups are associated with these proteins including 156 chlorophylls, 37 carotenoids, 2 phyloquinones, and 3 iron-sulfur clusters (4Fe-4S) (Caspy and Nelson 2018). During *in vivo* linear electron transport, an initial light-induced charge separation occurs between P_700_ (a chl a•chl *a’* dimer) and A_0_, forming P_700_^+^ and A_0_^−^, this is followed by sequential electron transfer from A_0_^−^ to A_1_ (phylloquinone), the iron-sulfur clusters F_X_ and F_A_/F_B_. Subsequently, oxidized ferredoxin is reduced on the stromal side of the thylakoid membrane and P_700_^+^ is reduced by plastocyanin on the lumenal side of the membrane. It has been shown that electron transfer from P_700_ to F_x_ in PS I-LHC I can occur down both pseudosymmetrical cofactor branches associated with the PsaA/PsaB core dimer (Guergova-Kuras et al. 2001).

The production of reactive oxygen species (ROS) is unavoidable, being a consequence of electron transport under oxic conditions being produced by both respiratory electron transport membrane protein complexes and those of oxygenic photosynthesis. ROS are generated by all of the photosynthetic membrane protein complexes (PS II, the *b*_*6*_/*f* complex, PS I and, probably, the chloroplast NAD(P) dehydrogenase-like complex) and these ROS species can oxidatively damage proteins and other biomolecules (Das and Roychoudhury 2014). ROS are formed by the excitation of molecular oxygen (singlet oxygen, ^1^O_2_), the partial reduction of molecular oxygen (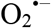), and the partial oxidation of water (H_2_O_2_ and 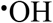). Additionally, dismutation of 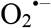 can form H_2_O_2_ that subsequently may be converted to 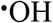 via Fenton chemistry. It has been estimated that ROS damage to PS II, alone, could account for a 10% loss in photosynthetic productivity (Long et al. 1994)). While ROS species are typically thought to be agents of damage, it should be noted that ROS also serve as important signaling molecules (Foyer and Noctor 2013; Mignolet-Spruyt et al. 2016).

ROS generation by PS II has been extensively studied (Kale et al. 2017; Pospíšil 2009; Pospíšil 2016), however, far fewer studies have been performed on PS I (Sonoike 2011). Under most laboratory conditions (constant moderate temperature and constant, relatively low intensity illumination) PS I appears to be significantly less susceptible to photodamage than PS II (Sonoike 2011). However, in the field environment, plants are exposed to a variety of abiotic stressors which can lead to enhanced ROS production (drought, low or high temperatures, highlight intensities, nutrient limitations, etc.) (Choudhry et al. 2016; You and Chan 2015). Under such field conditions, PS I can undergo significant photoinhibition (Sejima et al. 2014; Sonoike et al. 1995; Terashima et al. 1994; Zivcak et al. 2015).

In our study, we have used high-resolution tandem mass spectrometry to identify the location of oxidized residues within PS I-LHC I supercomplex isolated from field-grown spinach. Earlier, we have used these methods to identify natively oxidized residues in spinach PS II (Frankel et al. 2012; Frankel et al. 2013), results which have been largely verified and extended for cyanobacterial PS II (Weisz et al. 2017). We have also used these methods to examine natively oxidized residues in the cytochrome *b*_*6*_/*f* complex (Taylor et al. 2018).

In this communication we have mapped the identified natively oxidized residues in the spinach PSI-LHCI supercomplex and mapped these onto the corresponding residues of the *Pisum sativum* PSI-LHCI supercomplex structure (Mazor et al. 2017). We observed many oxidized residues on the surface of the complex. This is expected since ROS produced by any source, either on the stromal or lumenal sided of the thylakoid, could potentially modify surface residues. Additionally, a group of largely buried oxidized residues lead from the chl *a’* of the P_700_ special pair to the surface of the complex. Two buried oxidized residues were also in relatively close proximity to A_1B_. These findings support the hypothesis that the P_700_ and, possibly, A_1B_ are sources of ROS production by the PS I-LHC I supercomplex, probably ^1^O_2_ and 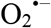, respectively. Finally, a large number of buried residues located in the hydrophobic cores of Lhca1-Lhca4, the distal antenna for PS I, are oxidatively modified. These appear to be the result of ^1^O_2_ production by the distal antennae for the photosystem. Interestingly, no oxidatively modified residues were identified in close proximity to the 4Fe-4S clusters F_X_, F_A_ or F_B_. These metal centers have been proposed to be either a source of 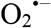 or a target for 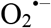 damage (Asada 1999; Caffarri et al. 2014; Tjus et al. 1999).

## Materials and Methods

PS I-LHC I-LHC II enriched membranes were prepared essentially as described previously (Bell et al. 2015). The PS I-LHC I-LHC II subunits were resolved by LiDS-PAGE (Delepelaire and Chua 1979), using a non-oxidizing gel system (Rabilloud et al. 1995). It is known that standard SDS-PAGE can introduce numerous protein oxidation artifacts (Sun and Anderson 2004) due to the generation of 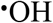 during the polymerization process. In this system, the acrylamide solution is degassed and the gels are polymerized with riboflavin (in the presence of diphenyliodonium chloride and toluensulfinate) with overnight exposure to long-wavelength UV light. During electrophoresis the upper reservoir buffer contained thioglycolate. It has been demonstrated that proteins resolved in this system exhibit significantly lower levels of electrophoresis-induced protein oxidation than proteins resolved by standard PAGE using standard polymerization conditions, confirming earlier reports (Frankel et al. 2012; Rabilloud et al. 1995; Sun and Anderson 2004). Electrophoresis was stopped when the proteins entered the resolving gel ~1.0 cm. The gel was then stained with Coomassie blue, destained, and two protein bands that contained, 1. principally PsaA/PsaB and, 2. The low molecular weight PS I core subunits and LHCs, were excised and analyzed separately. These were digested using either trypsin or chymotrypsin using standard procedures for “in-gel” proteolysis. After protease digestion and peptide isolation, the peptides were resolved by HPLC on a C:18 reversed phase column and ionized via electrospray into a Thermo Scientific Orbitrap Fusion Lumos mass spectrometer. The samples were analyzed in a data-dependent mode with one Orbitrap MS^1^ scan acquired simultaneously with up to ten linear ion trap MS^2^ scans. The MassMatrix Program (Xu and Freitas 2009) was used for protein identification and the identification of peptides containing oxidative modifications. A FASTA library containing the sequences of 19 proposed subunits of PS I-LHC I of the spinach supercomplex was searched, as was a decoy library containing the reversed amino acid sequences of the 19 proposed subunits. If a search exhibited any hits to this decoy library (i.e. decoy hit % > 0.00%) it was excluded from further analysis. It should be noted that for standard proteomic analysis in MassMatrix, a decoy hit rate of up to 10% is tolerated. Twelve different oxidative modifications were considered during peptide screening with a maximum of two modifications allowed per peptide. For a putative positive identification of an oxidized residue by MassMatrix, the peptide containing the modification must exhibit a *pp-tag* value of 10^−5^ or smaller and either *pp* or *pp2* must exhibit a value of 10^−5^ or smaller (Bricker et al. 2015; Xu and Freitas 2009). Peptides meeting these extremely rigorous *p*-value thresholds were then examined manually and the quality of the MS^2^, collision-induced dissociation spectra, being confirmed. It should be noted that typically, during peptide identification by mass spectrometry for proteomic experiments, *p*-values of 5 × 10^−2^ are typically used (Diz et al. 2011). We feel that this is inadequate when identifying posttranslational modifications (Bricker et al. 2015). Additionally, only peptides with charge states of ^+^3 or lower were considered. Finally, the mass error of the precursor ion was required to be ≤ 5.0 ppm and identified as the product of specific proteolytic cleavage. The identified oxidized amino acid residues were mapped onto the crystal structure of the *Pisum satvium* PS I-LHC I supercomplex (PDB: 5L8R, (Mazor et al. 2017)) using PYMOL (DeLano 2002).

## Results and Discussion

The isolation of the spinach PS I-LHC I-LHC II membranes was performed as previously described and the protein composition of these membranes (Fig. 1) was basically indistinguishable from our previous report (Bell et al. 2015). The PS I-LHC I-LHC II membranes were highly enriched in PS I core components and various light-harvesting chlorophyll proteins (Lhcas and Lhcbs) and were highly depleted of PS II core components. We had earlier shown that these membranes were also depleted of cytochrome b_6_/*f* and CF_1_-CF_o_ subunits (Bell et al. 2015). Additionally, we had demonstrated that the LHC II associated with PS I were functionally coupled to the photosystem (Boss et al. 2017) and that there was little or no LHC II which was not energetically coupled to the photosystem (Bell et al. 2015). All available evidence indicates that these membranes contain large quantities of PS I-LHC I supercomplexes that are functionally coupled to LHC II trimers.

**Figure 1.**
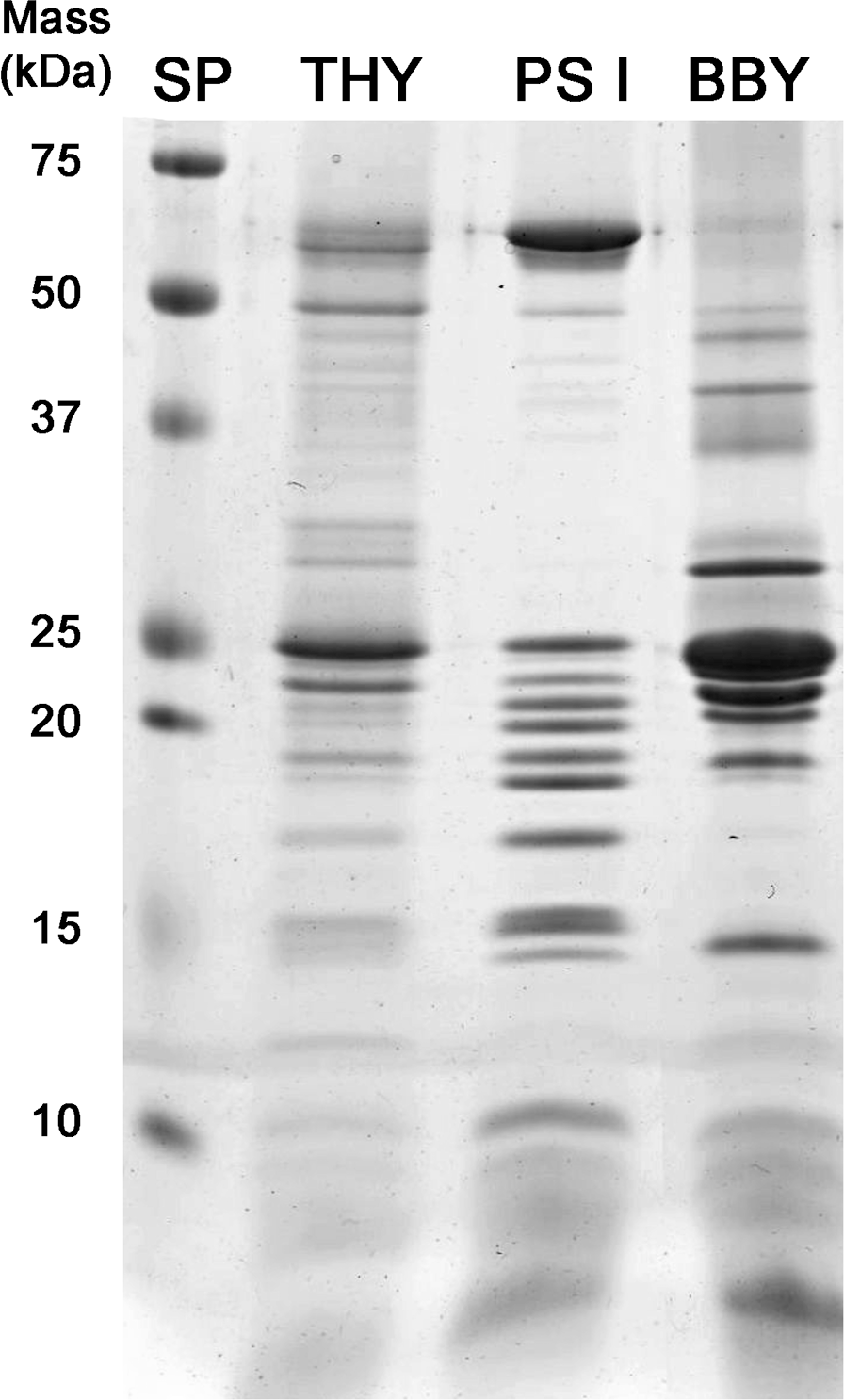
LiDS-PAGE of Thylakoids, PSI-LHCI-LHCII membranes, and PS II membranes. Individual lanes were loaded with 4 μg chl. The PSI-LHCI-LHCII membranes are highly enriched in PSI core proteins, contain significant amounts of LHCs (both lhcas and lhcbs), and are highly depleted of PS II components (BBY) as previously reported (Bell et al. 2015). THY, thylakoids; PSI, PSI-LHCI-LHCII membranes; BBY, PS II membranes. Standard proteins (SP) are shown to the left.

Tandem mass spectrometry analysis of the tryptic and chymotryptic peptides of the PS ILHC I supercomplex allowed the identification of 166 oxidatively modified residues present on the subunits of the supercomplex. In this study over 200,000 peptides were examined and screened for the presence of oxidative modifications. The mass spectrometry sequence coverage and the number of oxidatively modified residues observed for each PS I-LHC I subunit are shown in Table 1. One should note that most of the subunits of the PS I-LHC I supercomplex are intrinsic membrane subunits. Intrinsic membrane proteins are often difficult to examine by high resolution tandem mass spectrometry. Typically, only low sequence coverage being reported in standard experiments (Kar et al. 2017; Souda et al. 2011; Weisz et al. 2017). In our experiments, high sequence coverage was obtained for most of the subunits of the supercomplex, ranging from 100% for PsaA, PsaJ and Lhca4 to 56% for PsaG. The use of chymotrypsin greatly expanded the portfolio of observed peptides and dramatically increases the sequence coverage since the chymotryptic peptides are, in large measure, complementary in sequence to the tryptic peptides. Peptides were detected for PsaI, PsaK, PsaO and PsaP; however, the protein identification based on these peptides exhibited a significant number of hits to the decoy library. Since we set very rigorous criteria for protein identifications (i.e. 0.00% hits to the decoy library were allowed, see Materials and Methods), these proteins were excluded from subsequent analysis. No peptides were observed for PsaN. The number of oxidized amino acid residues observed was quite variable, ranging from 42 for PsaB (5.7% of the PsaB residues being modified) to 0 for PsaG (0% of the PsaG residues being modified) indicating that some supercomplex subunits are significantly more susceptible to oxidative modification than others.

**Table 1.**
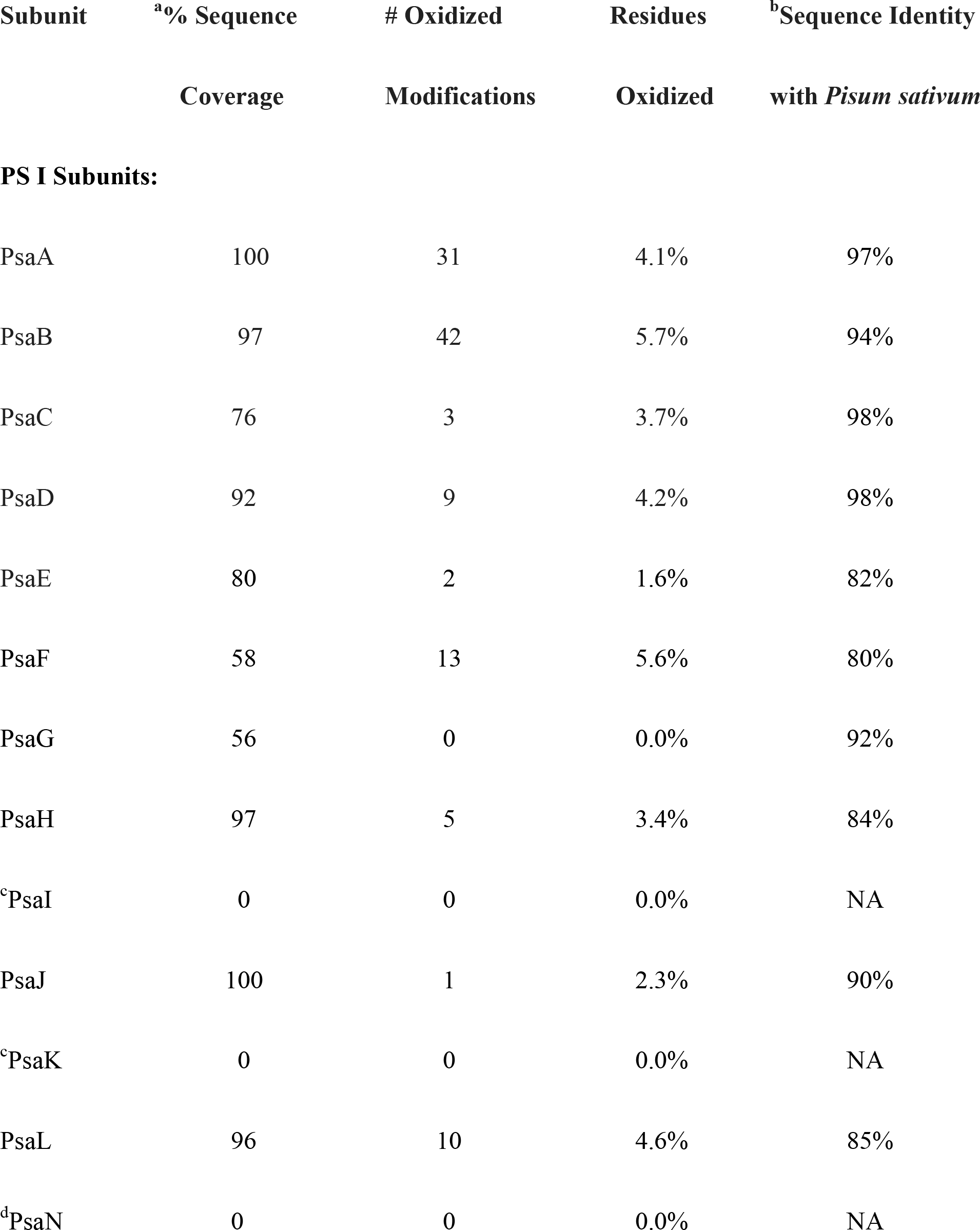
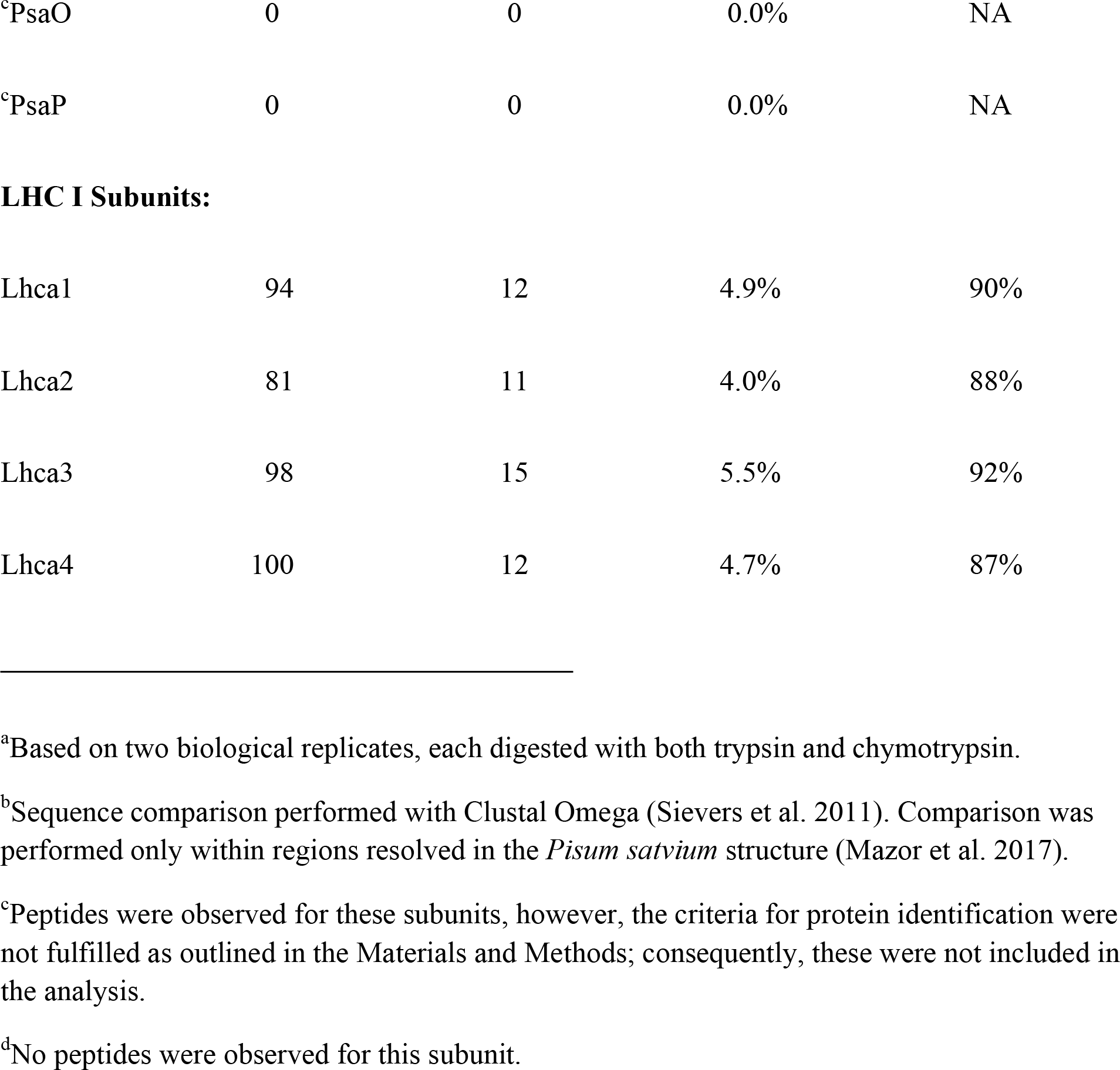
Summary Data for Mass Spectrometry and Sequence Comparison of Spinach PS I to *Pisum sativum* PS I.

| Subunit | <sup>a</sup>% Sequence Coverage | # Oxidized Modifications | Residues Oxidized | <sup>b</sup>Sequence Identity with *Pisum sativum* |
| --- | --- | --- | --- | --- |
| **PS I Subunits:** |  |  |  |  |
| PsaA | 100 | 31 | 4.1% | 97% |
| PsaB | 97 | 42 | 5.7% | 94% |
| PsaC | 76 | 3 | 3.7% | 98% |
| PsaD | 92 | 9 | 4.2% | 98% |
| PsaE | 80 | 2 | 1.6% | 82% |
| PsaF | 58 | 13 | 5.6% | 80% |
| PsaG | 56 | 0 | 0.0% | 92% |
| PsaH | 97 | 5 | 3.4% | 84% |
| <sup>c</sup>PsaI | 0 | 0 | 0.0% | NA |
| PsaJ | 100 | 1 | 2.3% | 90% |
| <sup>c</sup>PsaK | 0 | 0 | 0.0% | NA |
| PsaL | 96 | 10 | 4.6% | 85% |
| <sup>d</sup>PsaN | 0 | 0 | 0.0% | NA |
| <sup>c</sup>PsaO | 0 | 0 | 0.0% | NA |
| <sup>c</sup>PsaP | 0 | 0 | 0.0% | NA |
| **LHC I Subunits:** |  |  |  |  |
| Lhca1 | 94 | 12 | 4.9% | 90% |
| Lhca2 | 81 | 11 | 4.0% | 88% |
| Lhca3 | 98 | 15 | 5.5% | 92% |
| Lhca4 | 100 | 12 | 4.7% | 87% |

<sup>a</sup>Based on two biological replicates, each digested with both trypsin and chymotrypsin.
<sup>b</sup>Sequence comparison performed with Clustal Omega (Sievers et al. 2011). Comparison was performed only within regions resolved in the *Pisum satvium* structure (Mazor et al. 2017).
<sup>c</sup>Peptides were observed for these subunits, however, the criteria for protein identification were not fulfilled as outlined in the Materials and Methods; consequently, these were not included in the analysis.
<sup>d</sup>No peptides were observed for this subunit.

Fig. 2 shows the quality of the data used for the identification of oxidized amino acid residues within the PS I-LHC I supercomplex. The tandem mass spectrometry data collected for the ^303^D-^314^R tryptic peptide of PsaB is illustrated. This peptide bears a general oxidation modification (mass change ^+^16) at ^308^M. The observed mass accuracy for the parent ion was 2.6 ppm. The *pp*, *pp*_*2*_ and *pp*_*tag*_ values for this peptide were 10^−6.3^, 10^−6.3^ and 10^−5.5^ (Xu and Freitas 2009), respectively, and is consequently among the lowest quality peptides used in this study (Bricker et al. 2015). Even this peptide, however, exhibited nearly complete y- and b-ion series, allowing unequivocal identification and localization of the oxidative mass modification. This result indicates that the use of *p* values ≤ 10^−5^ provided high quality peptide identifications for the localization of post-translational modifications.

**Figure 2.**
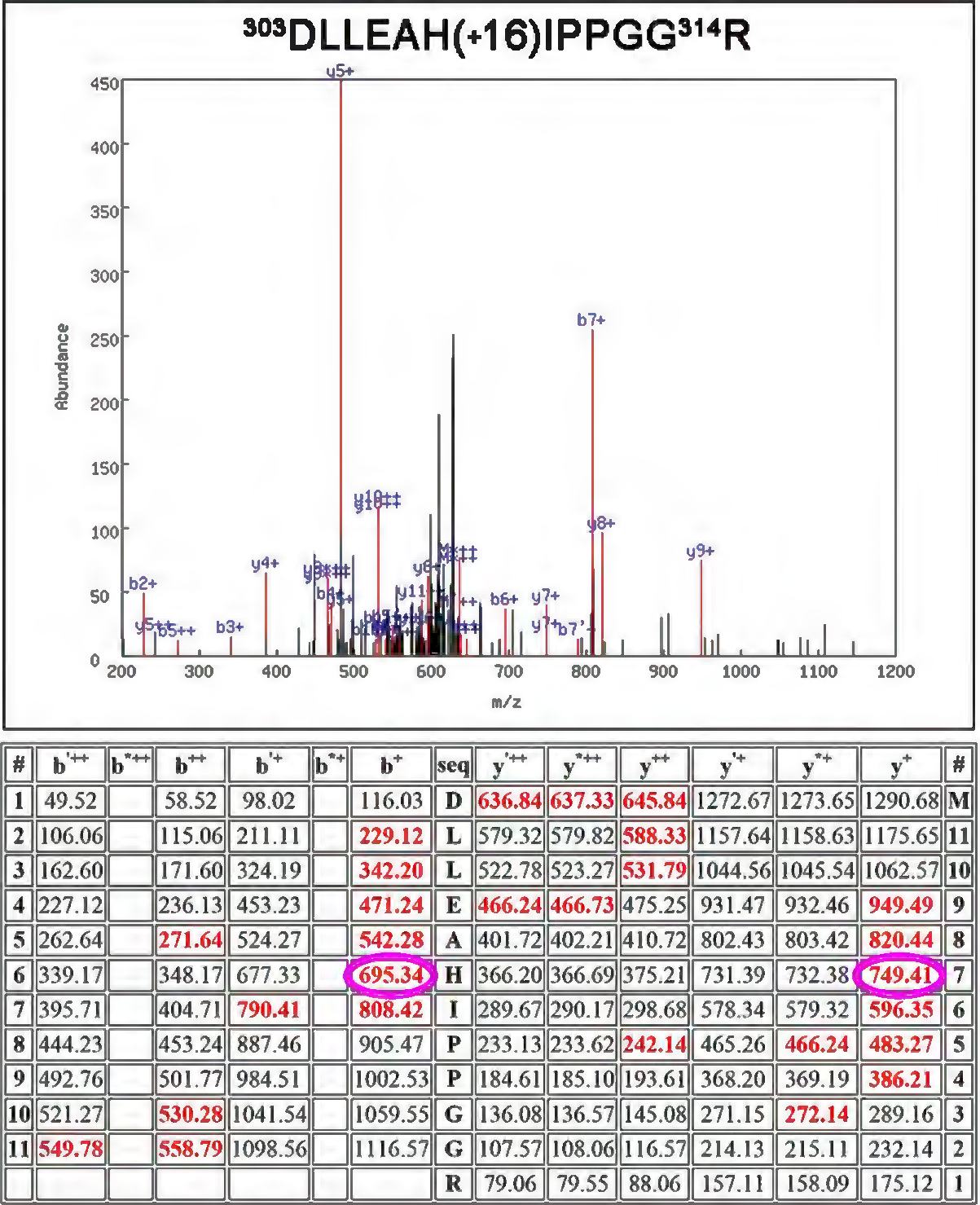
Quality of the Mass Spectrometry. Shown is the mass spectrometry result for the tryptic peptide PsaB: ^303^D-^314^R which contains a modified ^308^H + go (+16 amu) modification. Top, spectrum of the CID dissociation of the unmodified peptide PsaB:^303^DLLEAH (+16)IPPGG314R. Various identified ions are labeled. Bottom, table of all predicted masses for the y- and b-ions generated from this peptide sequence. Ions identified in the CID spectrum (top) are shown in red. The b^’++^, b^’+^ y^’++^ and y^’+^ ions are generated by the neutral loss of water while the b^*++^, b^*+^ y^*++^ and y^*+^ ions are generated from the loss of ammonia. For our analysis, only y^X^ and b^X^ ions were considered. The ions y7^+^-y9^+^ and y10^++^-y12^++^ exhibit the +16 mass modification as does the b6^+^-b7^+^ and b10^++^-b11^++^ ion. This verifies that 308H contains an oxidative modification which adds 16 amu to the histidyl residue. The *p*-values for this peptide were *pp* = 10^−6.3^, *pp*_*2*_ = 10^−6.3^ and *pp*_*tag*_ = 10^−5.5^.

The identity of these oxidized residues and the types of modifications observed are presented in Table 2. It should be noted that it is extremely unlikely that all of the observed modifications would be present on every copy of the complex. ROS modification of a particular amino acid residue is a stochastic rather than mechanistic process. The probability that a residue will be modified depends on the ROS type, the susceptibility of a residue to a particular ROS type, the residency time of the ROS in proximity to the residue, and the lifetime of the ROS species. Our working hypothesis is that residues located near a site of ROS production (or along an ROS exit pathway) have a higher probability of oxidative modification than those distant from a ROS production site (or not along an ROS exit pathway). Consequently, these detectable modifications are present within the full population of PS I-LHC I complexes present in our biological samples. The supercomplex was isolated from field-grown market spinach, consequently, the exact growth parameters are unknown. What is virtually certain, however, is that the plant material used in our study has been exposed to biologically relevant stress conditions (transient drought, low/high temperatures, high light intensities, fluctuating light intensity, nutrient limitations, etc.) typical of plants grown in the field (Choudhry et al. 2016; You and Chan 2015).

**Table 2.**
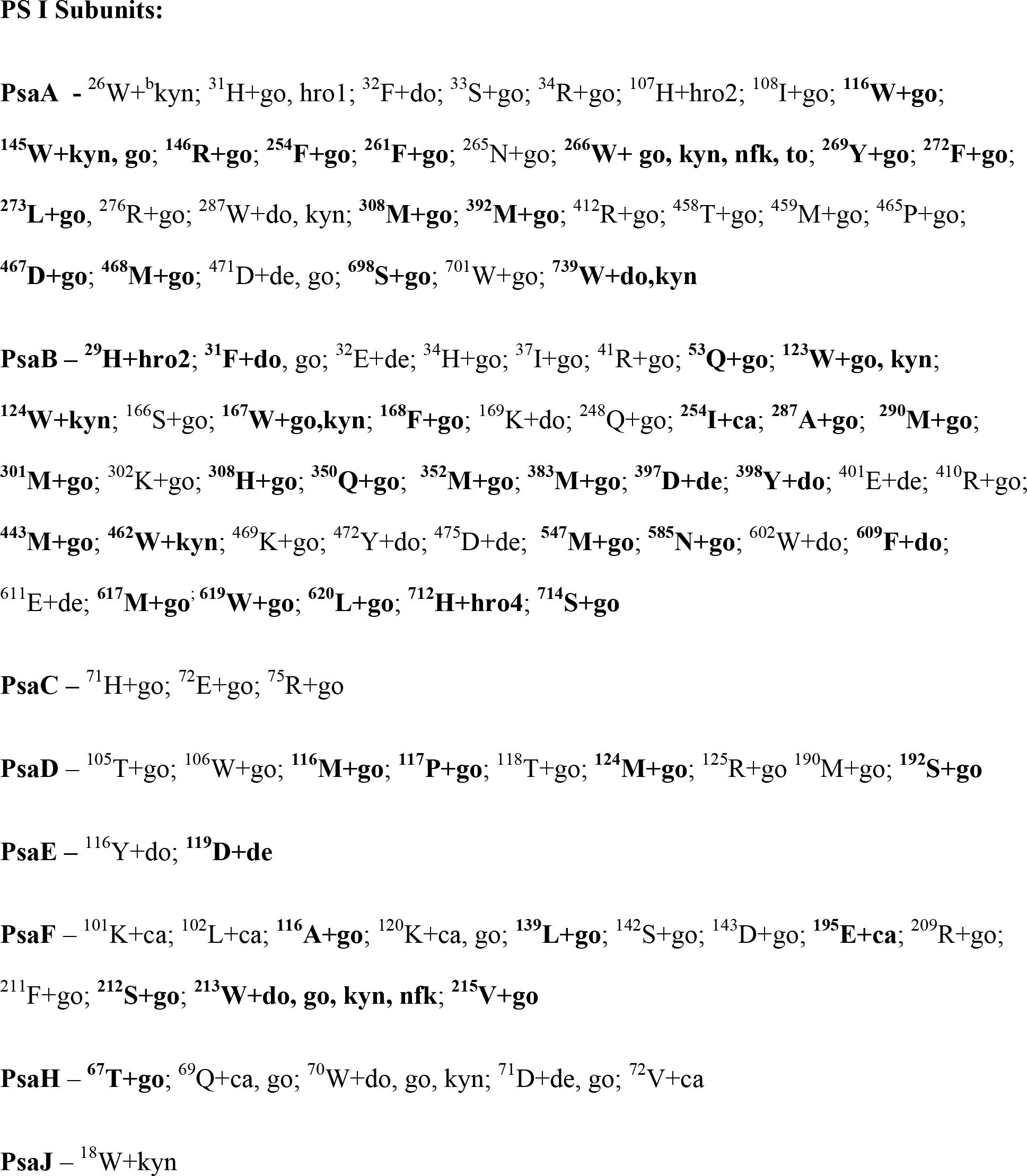
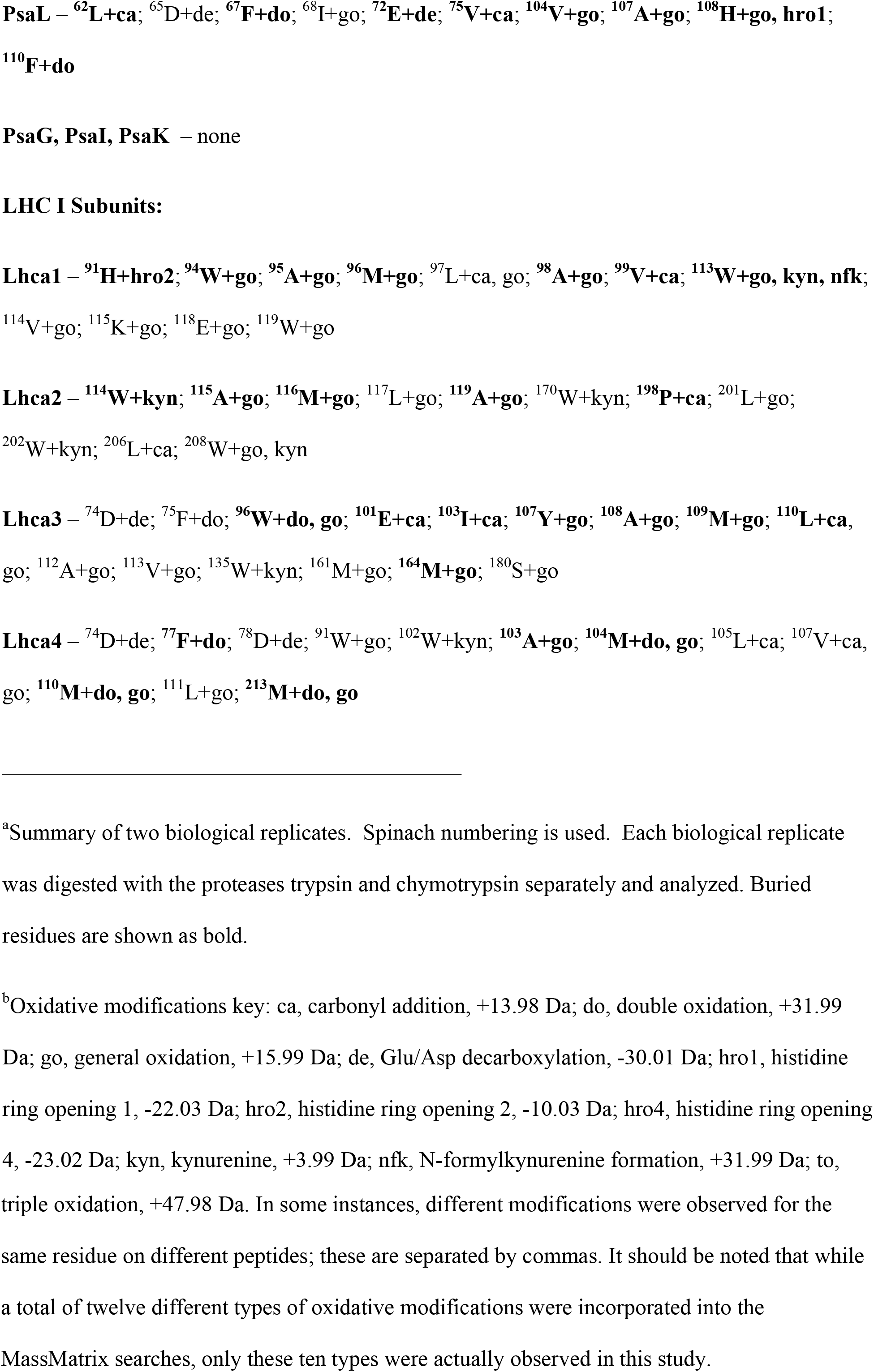
^a^Natively Oxidized Residues in the PS I-LHC I Supercomplex

No high-resolution structure is available for the spinach PS I-LHC I supercomplex. However, structures are available for a variety of other eukaryotic organisms: the red alga *Cyanidioschyzon merolae*, (Antoshvili et al.); the green alga *Chlamydomonas reinhardtii*, (Su et al. 2019; Suga et al. 2019); the moss *Physcomitrella patens* (Iwai et al. 2018) and the angiosperm *Pisum satvium* (PDB: 5L8R, (Mazor et al. 2017)). The sequence identity between the spinach subunits and the Pisum subunits is quite high, ranging from 98% for PsaC and PsaD to 80% for PsaF (Table I). This high degree of identity allowed us to map the oxidized amino acids that we observed in the spinach supercomplex onto the crystal structure of the *Pisum* supercomplex with 94% of the observed oxidatively modified residues being identical in both spinach and *Pisum*.

Fig. 3 presents a broad overview of the location of the oxidized residues that we identified within the context of the PS I-LHC I supercomplex. 49% of the oxidized residues were surface localized. This was not unexpected, as surface domains are particularly susceptible to oxidative modification. ROS produced by a variety of different sources including PS II (HO^.^, H_2_O_2_, ^1^O_2_ and 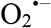), the cytochrome b_6_ *f* complex (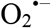 and ^1^O_2_), and PS I, itself (^1^O_2_, 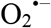 and, indirectly, H_2_O_2_ and HO^.^) are released to the stromal and lumenal compartments and may contribute to the oxidative modification of surface-exposed domains of the PS I-LHC I supercomplex, the exposed domains of other membrane protein complexes, and soluble proteins. Many ROS detoxification systems present in both the stromal (Das and Roychoudhury 2014; Tripathy and Oelmuller 2012) and lumenal compartments (Bermudez et al. 2012; Levesque-Tremblay et al. 2009). However, it is highly unlikely, given the possibly large amounts of ROS produced during photosynthetic electron transport, that these that these would offer complete protection. Additionally, the ROS detoxification systems would be ineffective against any ROS produced within the protein matrix (at least until the ROS was released to the bulk solvent). Indeed, we observe numerous oxidized residues (51%) buried within the protein matrix. We hypothesized that a subset of these modified residues would be in close proximity to the site(s) of ROS production within the photosystem,

^1^O_2_ has been implicated in PS I photoinhibition in PS I submembrane preparations (Rajagopal et al. 2005), *in vitro*, and under repetitive light pulse conditions, in vivo (Takagi et al. 2016). Within the photosystem several cofactors have been proposed to be the sources of this ROS. These include ^3^P_700_ (Rutherford et al. 2012) and the antennae chls (Alboresi et al. 2009; Cazzaniga et al. 2012; Krieger-Liszkay 2004) as possible sources of ^1^O_2_. It has also been shown that 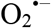 contributed to PS I photoinhibition under both chilling (Sonoike and Terashima 1994; Sonoike et al. 1995; Terashima et al. 1994) and repetitive light pulse conditions (Takagi et al. 2016). Additionally, dismutation of 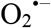 by superoxide dismutase leads to the formation of H_2_O_2_ which, in the presence of reduced iron-sulfur clusters or free iron, may generate the highly reactive HO^.^ (Sonoike et al. 1995; Takahashi and Asada 1988) via Fenton chemistry. The sources of this ROS are under some debate. The phylloquinone A_1B_ (Takagi et al. 2016) and the iron-sulfur clusters F_X_, F_A_ and F_B_ have been suggested as a possible source of 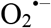(Asada 1999; Tjus et al. 1999).

**Figure 3.**
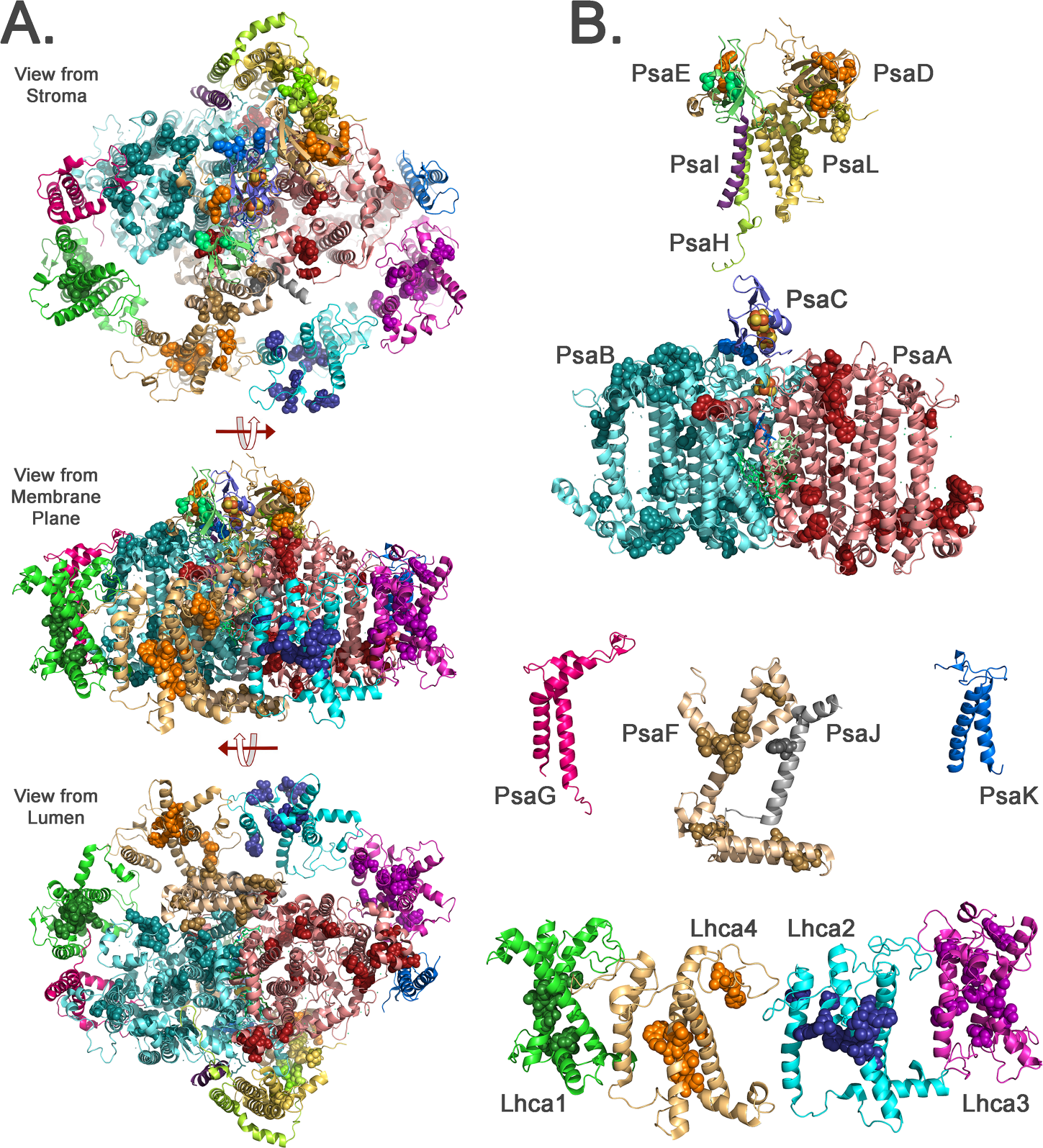
Overview of Natively Oxidized Amino Acid Residues in the Spinach PS I - LHC I Supercomplex. A. Illustration of the entire supercomplex as viewed from the stroma (top), as viewed within the plane of the membrane (middle), and as viewed from the lumen (bottom). B. Exploded view of the supercomplex as viewed within the membrane plane. Individual subunits are color-coded and labeled in B. Oxidatively modified residues are shown as spheres and are colored in darker shades for each subunit. The observed oxidatively modified residues in spinach were mapped onto their corresponding locations on the *Pisum sativum* PS I - LHC I supercomplex structure (PDB: 5L8R, (Mazor et al. 2017)).

The core subunits of PS I, PsaA, PsaB and PsaC, contain all of the redox-active cofactors of the photosystem (Srinivasan and Golbeck 2009). These include: P_700_, the primary electron donor, which is a chl *a*.chl *a’* dimer (the chl a is bound to PsaB and the chl *a’* is bound to PsaA); A_A_ and A_B_, two “accessory” chl a bound to PsaA and PsaB, respectively; A_0A_ and A_0B_, chl a monomers bound to PsaA and PsaB, respectively, which serve as primary electron acceptors from P_700_; A_1A_ and A_1B_, phylloquinones bound to PsaA and PsaB, respectively; F_X_, a 4Fe-4S iron-sulfur cluster bound to both PsaA and PsaB; and F_A_ and F_B_, both 4Fe-4S iron-sulfur clusters bound to PsaC. These cofactors are arranged in two pseudosymetric branches along the PsaA/PsaB axis with electron transfer occurring down either branch (Guergova-Kuras et al. 2001). A number of oxidatively modified residues are present closely associated with these cofactors, as illustrated in Figs. 4 and 5. It should also be noted that PsaB exhibited a greater degree of oxidative modification than PsaA. 5.7% of the PsaB residues were modified whereas 4.2% of PsaA residues were modified (Table 1). This greater degree of oxidative modification of PsaB is fully consistant with earlier reports that PsaB appears more susceptable to oxidative damage, leading to its preferential loss during the PS I photoinhibition timecourse (Sonoike 1995; Sonoike 1996; Sonoike et al. 1997).

**Figure 4.**
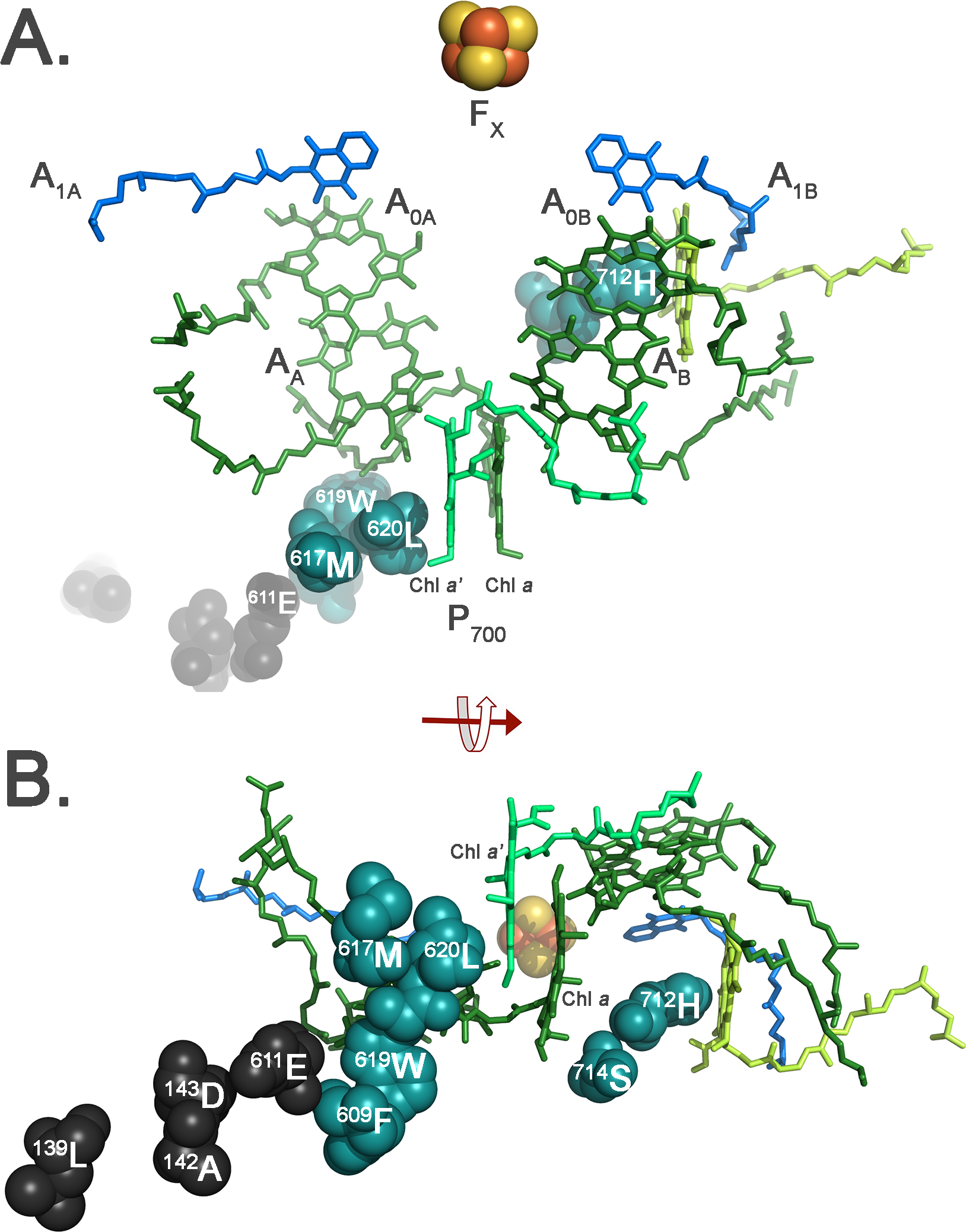
Details of the Oxidatively Modified Residues Identified in the Vicinity of the Cofactors Bound to PsaA and PsaB. A. View within the plane of the membrane. B. View from the lumenal face of the membrane. Cofactors are shown as sticks and are individually labeled. Oxidatively modified residues are shown as spheres. The residues PsaB: ^609^F, ^617^M, ^619^W, ^620^L, ^712^H, and ^714^S are buried and shown in blue. The residues PsaB:^611^E and PsaF: ^139^L, ^142^A and ^143^D are surface-exposed and shown in black. These residues form a near continuous path of modified residues leading from the chl a’ of P_700_ to the surface of the supercomplex. Please note that no PsaA residues were observed to be oxidatively modified in the vicinity of these cofactors. Additionally, the residue PsaB: ^712^H (and ^714^S) is in relatively close proximity to A_0B_ (8.2 Å), A1B (7.0 Å) and in close proximity to chl a 839 (4.6 Å), which is the closest chl to A_1B_.

**Figure 5.**
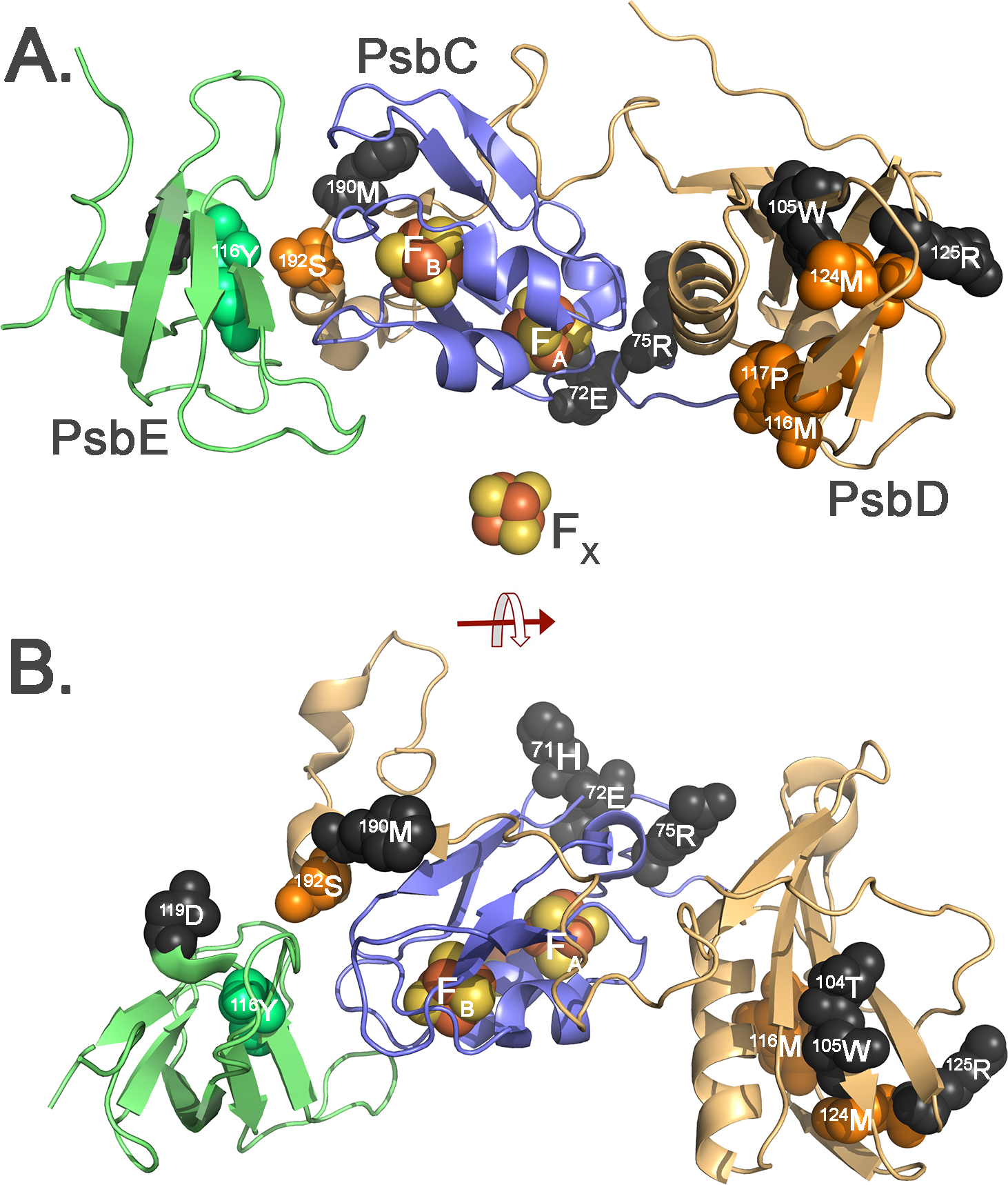
Details of the Oxidatively Modified Residues Identified in PsaC, PsaD and PsaE. A. View within the plane of the membrane. B. View from the lumenal face of the membrane. The three iron-sulfur clusters, F_X_, F_A_ and F_B_, are shown as spheres with F_X_ being included for orientation purposes. Oxidatively modified residues are shown as spheres. The residues PsaD: ^116^M, ^117^P, ^124^M and ^192^S and PsaE:^116^Y are buried and shown as orange spheres and green spheres, respectively. The residues PsaC: ^71^H, ^72^E and ^75^R, PsaD:^104^T, ^105^W, ^125^R and ^190^M, and PsaE:^119^D are all surface-exposed residues and are shown as black spheres. PsaC: ^75^R is relatively close to F_A_ (8.0 Å). All of the other oxidized residues present in these three subunits are >10 Å from the iron-sulfur clusters.

Figs. 4A and 4B illustrate the oxidized residues identified to be in close proximity to the redox-active cofactors associated with PsaA and PsaB. No oxidized residues were found in close proximity to F_x_; the nearest, PsaB:^547^M, is 10.6 Å distant. PsaB:^712^H and ^714^S are in relatively close proximity to A_1B_, with ^712^H (5.9 Å from A_1B_) being coordinated to chl *a* 839, which spacially is the closest chl to A_1B_. While this observation is consistant with the hypothesis that A_1B_ is a source of ROS, it should be noted that the chlorin ring of chl a 839 lies between A_1B_ and these oxidized residues. This complicates the interpretation, as the ROS responsible for the modification of PsaB:^712^H and ^714^S could have been generated at A_1B_, chl a 839, or other location.

Most interestingly, a chain of eight oxidized residues (PsaB:^620^L, ^617^M, ^619^W, ^609^F and ^611^E and PsaF: ^143^D, ^142^A, ^139^L) lead from the chl *a’* of P_700_ to the lumenal surface of the complex. No similar distribution of oxidized residues was identified associated with the chl a of P_700_. This does not appear to be due to sampling error since 100% mass spectrometry coverage was obtained for both PsaA and PsaB residues in this region. This chain of oxidized residues may define a prefered ROS exit pathway. Several studies have suggested that the triplet state of P_700_ (^3^P_700_) is preferentially localized to the chl *a’* (Breton et al. 1999; Li et al. 2004; Rutherford and Setif 1990). Importantly, the nearest carotenoid to the chl *a’* is nearly 11 Å distant and it is unlikley to efficiently quench ^3^P_700_. The interaction of ^3^P_700_ with dioxygen may lead to the formation of ^1^O_2_, and this may be a source of this ROS in the photosystem. Consequently, within the core of PS I, the production of ^1^O_2_ may center on the chl *a’* of P_700_. The presence of a defined set of oxidized residues leading away from the chl *a’* to to surface of the complex strongly supports this hypothesis.

Fig. 5 illustrates the locations of oxidatively modified amino acid residues in the vicinity of F_X_, F_A_ and F_B_. Interestingly, no residues were found in the immediate vicinity of any of these iron-sulfur clusters. Specifically, the nearest modified residue to F_X_ was PsaB:^547^M (10.6 Å), the nearest modified residue to F_B_ was PsaD:^192^S (11.6 Å) and the nearest modified residue to F_A_ was PsaC:^75^R (8.0 Å). The absence of proximal oxidative modifications associated with these cofactors is intriguing since the iron-sulfur clusters had been implicated in the formation of 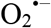 and possibly HO^.^ from H_2_O_2_ (Sonoike et al. 1995; Takahashi and Asada 1988). Damage to the 4Fe-4S clusters is also recognized as an early event during PS I photoinhibition (Sonoike et al. 1995; Takahashi and Asada 1988; Tjus et al. 1998). While F_x_ is buried within the PsaA/PsaB protein matrix and would be expected to have only limited accessibility to either 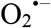 or H_2_O_2_, F_B_ is much more exposed, being 3-4 Å from the surface of PsaC. Interestingly, the PsaD and PsaE components exhibited a significant number of oxidative modifications. While the majority of these were localized on the surface of the complex, several were buried and proximally located near PsaC (PsaD: ^192^S, ^116^M, ^124^M; PsaE:^116^Y). Given the close proximity of PsaD and PsaE to PsaC, a working hypothesis is that ROS produced at the redox-active center(s) F_B_, and possibly F_A_, lead to the oxidative modifications observed on PsaD and PsaE. So the question arises, “why are no oxidative modifications observed in close proximity to F_B_ or F_A_ in the PsaC subunit?” Oxidative modifications may accumulate in the majority of PS I subunits with relatively limited effects on PS I function. If, however, oxidative damage occurs to F_B_ or F_A_ (or F_X_), electron flow to ferredoxin would be abolished, possibly triggering turnover and replacement of the entire PS I complex. This is the generally accepted model for PS I photoinhibition. Interestingly, it has been shown that under in vivo photoinhibitory conditions, PsaC was lost much more rapidly than other PS I core components in *Brassica* (Jiao et al. 2004), presumably due to damage of F_A_ and F_B_ (Sonoike 1996; Sonoike et al. 1995). In this study plants were grown under conditions similar to what the plants would experience in the field. A similar result was observed *in vitro* in spinach (Yu et al. 2000), with PsaC being among the most labile subunits during a photoinhibition timecourse (along with Lhca1-Lhca4, see below). This raises the intriguing possibility that under some conditions, damage to F_A_ or F_B_ leads to a more selective loss (and possibly replacement of PsaC). It should be noted that, in this hypothetical scenario, loss of PsaC would also lead to the release of PsaD and PsaE since PsaC is required for the binding of these subunits (at least in cyanobacteria (Yu et al. 1995)). If newly synthesized PsaC (or possibly PsaC bearing repaired 4Fe-4S clusters (Djaman et al. 2004)) was incorporated into PS I, released PsaD and PsaE, possibly bearing oxidized modifications, could be reincorporated into the photosystem.

Fig. 6 illustrates the locations of oxidative modifications in the Lhca1-Lhca4 subunits. These components serve as a distal antenna for PS I, delivering excitation energy to the core antenna which is associated with the PsaA and PsaB subunits. All four of the Lhcas exhibit significant oxidative modification. While many of the oxidatively modified residues are surface-exposed, many are buried near the hydrophobic cores of these subunits, along the helix A and B axis (Fig. 6A). In these locations the modified residues are in very close proximity to the lightharvesting chls and carotenoids (Fig. 6B).

**Figure 6.**
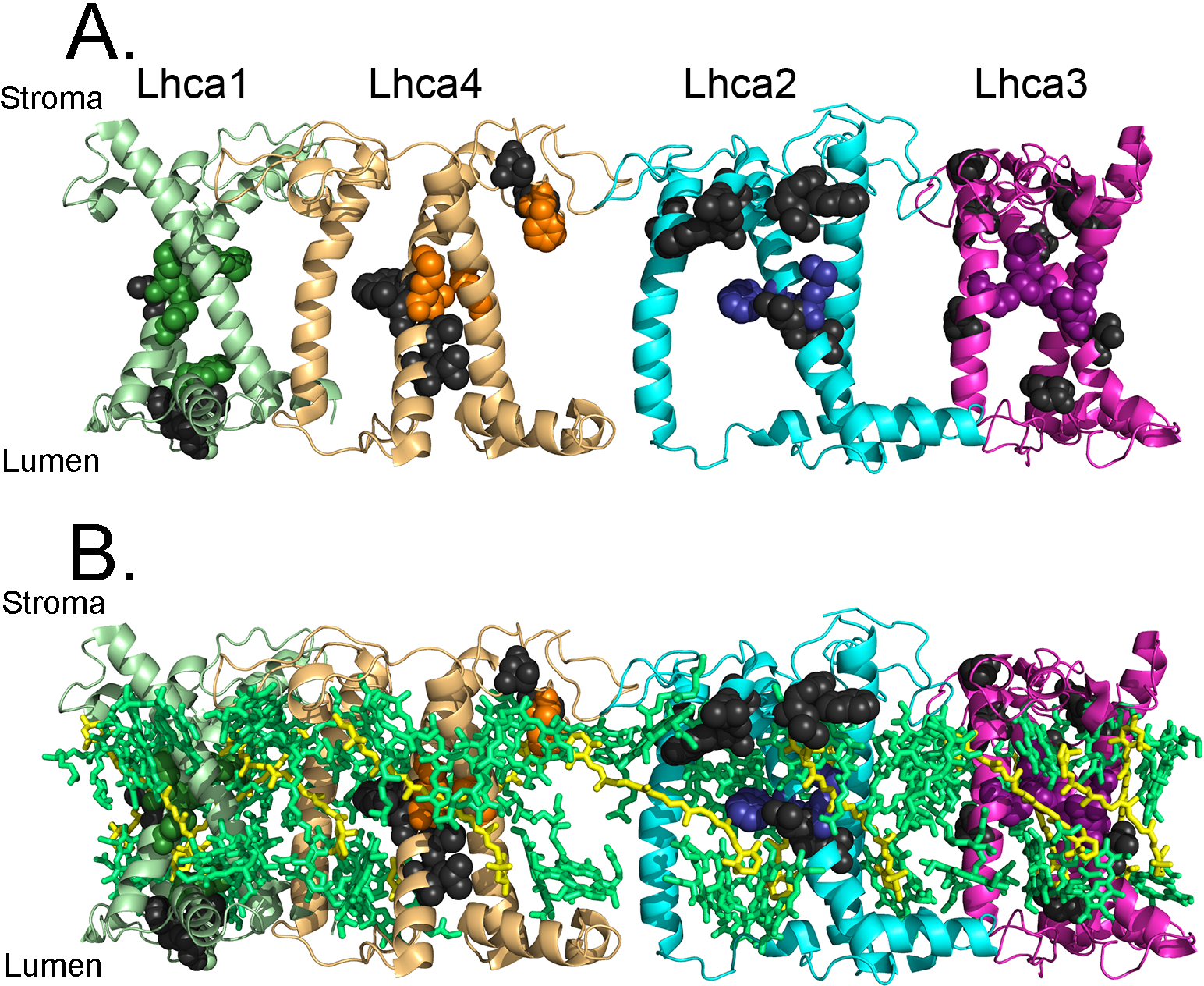
Modified residues in the Lhca1-Lhca4 distal antenna viewed within the plane of the membrane. A. Without cofactors shown. B. With cofactors shown. In A., individual Lhca subunits are labeled, as is the stromal and lumenal sides of the membrane complex. Modified residues are shown as spheres, with colored residues being buried and black residues being surface-exposed. Cofactors are shown in stick representation with chl (both chl a and chl b) shown in pale green and the various carotenoids, β-carotene, lutein, and violaxanthin, shown in pale yellow.

Earlier studies have indicated that the Lhcas are rapidly lost during PS I photoinhibition (Alboresi et al. 2009; Yu et al. 2000). The Lhca3 and Lhca4 contain “red” chl forms (Croce and van Amerongen 2013; Croce et al. 1996; Melkozernov and Blankenship 2005) with absorption bands of longer wavelengths than exhibited by P_700_. Excitation energy is concentrated in these low energy sinks prior to transfer to the proximal antenna and the reaction center. This slows excitation energy transfer; consequently there is a higher probability of intersystem crossing and formation of ^3^chl, and subsequently ^1^O_2_, at these sites. It has been suggested that the Lhca antenna acts as a “fuse” for the photosystem (Alboresi et al. 2009). Under conditions which promote PS I photoinhibition (high light intensity and acceptor-side electron transport limitation), the Lhca antenna chlorophyll are degraded in parallel with the Lhca proteins, while the reaction center retains photochemical activity (Alboresi et al. 2009). In this scenario, any ^3^chl which is formed is directly quenched by a proximal carotenoid, if this fails and ^1^O_2_ is generated, then this is quenched either physically or chemically by the carotenoid, if this fails ^1^O_2_ either escapes the Lhca or oxidatively damages this component (either the Lhca protein or its chl cofactors), perhaps leading to Lhca turnover. This would not have a effect on PS I mechanistically, but would directly lead to a smaller optical cross-section for the photosystem. Ultimately, this would offer some degree of protection for the photosystem from a high photon flux and/or an acceptor-side limitation. Our observation that the Lhca subunits exhibit significant oxidative modification strongly supports the hypothesis that these components are a source of ^1^O_2_ in the PS I-LHC II supercomplex.

## Conclusions

In this communication, we have identified oxidized residues in the PS I- LHC I supercomplex. The locations of these support our hypothesis that residues in the vicinity of ROS production sites would be particularly prone to oxidative modification. A clearly defined group of oxidatively modified residues appears to lead from the chl *a’* of P_700_ to the surface of the complex. This result supports the hypothesis that ^3^P_700_, which is localized to the chl *a’*, participates in the formation of ^1^O_2_. Two oxidized residues are located in proximity to A_1B_ (PsaB:^712^H and ^714^S). This finding is consistent with the hypothesis that A_1B_ may be a source of ROS, probably 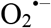. Additionally, numerous oxidized residues are located in Lhca1-Lhca4. Many of these are buried in the hydrophobic domains of these subunits and are in close proximity to the chls and carotenoid cofactors. This observation is consistent with the hypothesis that the Lhcas are a source of ^1^O_2_. Finally, no oxidized residues were observed in close proximity to the iron-sulfur clusters F_X_, F_A_ or F_B_. This observation indicates that oxidative damage associated with these cofactors, which would interrupt electron flow to ferredoxin, may trigger turnover of the entire PS I-LHC I supercomplex or, possibly a repair pathway leading to the insertion of undamaged or repaired PsaC. We have not, at this time, determined the relative importance of the various possible sites of ROS production for photoinhibition of PS I, nor the time course for the appearance of oxidative modifications during the photoinhibition process. These important questions are the subject of future studies.

## Acknowledgements

This work was solely supported by the United States Department of Energy, Office of Basic Energy Sciences grant DE-FG02-09ER20310 to TMB and LKF.

